# Unveiling the three-step model for the interaction of imidazolium-based ionic liquids on albumin

**DOI:** 10.1101/2023.05.25.542168

**Authors:** Juliana Raw, Leandro R. Franco, Luiz Fernando de C. Rodrigues, Leandro R. S. Barbosa

## Abstract

The effect of the ionic liquids (ILs) 1-methyl-3-tetradecyl imidazolium chloride ([C_14_MIM][Cl]), 1-dodecyl-3-methylimidazolium chloride ([C_12_MIM][Cl]) and 1-decyl-methylimidazolium chloride ([C_10_MIM][Cl]) on the structure of bovine serum albumin (BSA) was investigated by fluorescence spectroscopic, UV-Vis spectroscopy, small angle X-ray scattering and molecular dynamics simulations. Concerning the fluorescence measurements, we observed a blue shift and a fluorescence quenching as IL concentration increased in the solution. Such behavior was observed for all three studied imidazolium-based IL, being larger as the number of methylene groups in the alkyl chain grew. UV-Vis absorbance measurements indicate that even at relatively small IL:protein ratios, like 1:1, or 1:2 ([C_14_MIM][Cl]) is able to change, at least partially, the sample turbidity. SAXS results agree with the spectroscopic techniques and suggest that the proteins underwent a partial unfolding, evidenced by an increase in the radius of gyration (*R_g_*) of the scattering particle. In the absence and presence of ([C_14_MIM][Cl])=3mM BSA *R_g_*, increases from 29.1 to 45.1 Å, respectively. Together, these results indicate that the interaction of BSA with IL is divided into three stages: the first stage is characterized by the protein in its native form. It takes place for IL:protein *≤* 1:2 and the interaction is predominantly due to the electrostatic forces, provided by the negative charges on the surface of the BSA and the cationic polar head of the ILs. In the second stage, higher IL concentrations induce the unfolding of the protein, most likely inducing the unfolding of domains I and III, in such a way that the protein’s secondary structure is kept almost unaltered. In the last stage, IL micelles start to form and, therefore, interaction with protein reaches a saturation point and free micelles may be formed. We believe this work provides new information about the interaction of ILs with BSA.

**Graphical Abstract:** 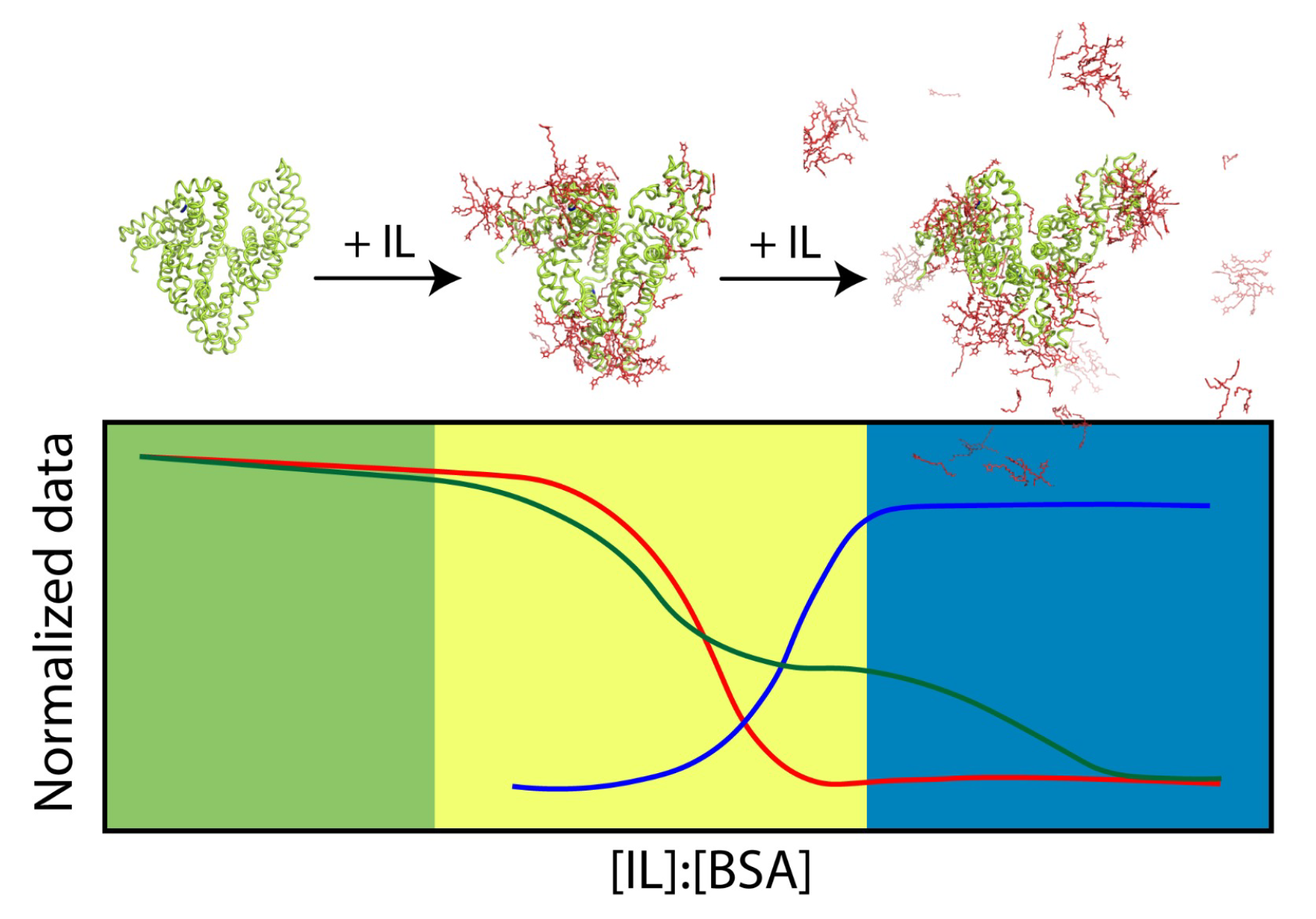

## Introduction

The development of sustainable procedures and the use of molecules with smaller harmfulness to the environment has been encouraged by the awareness of the industry and regulatory agencies. Organic solvents are one of the main environmental impact factors in chemical processes, so greener substitutes are increasingly in demand. Ionic liquids (ILs) are the result of these efforts and, in particular, surfactant ionic liquids have shown great success in this area.^1^ ILs are salts that are liquid at temperatures smaller than 100°C. ILs have interesting physicochemical properties like negligible vapor pressure, low degree of flammability, and high ionic conductivity. ^2, 3^ The small vapor pressure ensures the ionic liquids its “green chemistry” use.^4^ Since ILs can be exploited as an alternative solvent to the traditional organic ones (like chloroform or hexane), they can be used to improve electronic components^5, 6^ and also in biocatalysis processes.^7, 8^

Protein-surfactant interactions are an important field of study due to their applications in personal care, pharmaceuticals, cosmetics, detergents, and bioscience.^9, 10^ Surfactants can denature,^11^ unfold^12^ and refold proteins,^9, 10, 13^ according to the properties, structure, and concentration of the protein and the surfactant being studied.^14^ The nature of both molecules also determines the predominant interaction between the three main forces of this system: electrostatic, hydrophobic, and Van der Waals.^15^ Despite a large amount of research on the interaction of ILs and surfactants with biologically relevant systems, a systematic study on imidazolium-based IL on model proteins, such as bovine serum albumin, is still indispensable.

Serum Albumin proteins are essential to life: they are considered transport proteins and are the most abundant protein in the blood at a concentration of *≈* 42*mg/ml*,^16^ with such high concentration being crucial to its stability. ^17^ Bovine Serum Albumin (BSA) has been used as a model protein for many biophysical and biochemical studies as a model protein; in particular for UV-Vis, steady-state fluorescence, and small-angle X-ray scattering (SAXS), due to their water solubility and flexible binding capacity and capability to be used at different pHs,^17^ urea concentrations^18^ and temperatures. BSA is made up of around 585 amino acid residues with a molecular weight of *≈* 66*kDa* and is negatively charged at physiological pH (since its pI is around 5.4^17^). From the spectroscopic point of view, BSA has two tryptophan residues (Trp134 and Trp212), and, when folded, Trp134 is located on the surface, more exposed to the solvent, and Trp212 is buried inside, less accessible. The interaction of model proteins with cationic amphiphilic molecules was already investigated in the literature: for instance, Chakraborty et al.^19^ studied the interaction of BSA with cationic surfactants like DTAB, TTAB and CTAB, non-ionic C_12_E(8) and anionic SDS surfactants. The authors reported that the BSA-SDS interaction is bimodal, while the BSA-cationic surfactants interaction increases with increasing tail length and is monomodal, according to the authors, cationic surfactants denatured BSA in a single step. ILs are expected to contribute in applications for drug delivery systems, but a better understanding of the mechanisms of action is needed.^20, 21^ This work provides new knowledge of the interaction between imidazolium-based IL and proteins. In light of this new information, the application of ionic liquids in systems of biological interest or the development of new ionic liquids may be more specific and accurate.

Herein, we studied the interaction of three alkyl-functionalized imidazolium-based ILs with BSA, in order to get more information on the effects of IL on different BSA structural levels. To do so, we used a combination of steady-state fluorescence, circular dichroism, small-angle X-ray scattering techniques, and molecular dynamics simulations.

## Materials and Methods

### Materials

Bovine Serum Albumin with stated purities higher than 99% was purchased from Sigma-Aldrich, Germany. 1-Methyl-3-tetradecylimidazolium chloride ([C_14_MIM][Cl]), 1-dodecyl-3-methylimidazolium chloride ([C_12_MIM][Cl]) and 1,3-didecyl-2-methylimidazolium chloride ([C_10_MIM][Cl]) with purity *>*98% are purchased from io-li-tec (Salzstrasse, Germany). All solutions were prepared in 20 mM acetate borate phosphate buffer (pH 7.3). Such buffer was chosen because we do know BSA behavior at pHs from 2.0 – 9.0.^17^ BSA solutions were freshly prepared and all measurements were performed at 22*±*1*^◦^*C. Stock so-lutions of Ils at 200mM were prepared at the same buffer. After this, samples were produced at the desired molar ratio among BSA and the ionic liquids.

## Methods

### Steady-state Fluorescence

Fluorescence measurements were performed using a Cary Eclipse – Varian spectrometer and a quartz cuvette of 2 mm optical path length. The fluorescence emission spectra were collected from 305nm to 500nm using an excitation wavelength of 295nm (i.e., using tryptophan as an intrinsic probe) and slit width for excitation of 5nm and for emission of 5nm. We chose 295 nm for the excitation wavelength to minimize the excitation of phenylalanine and tyrosine residues. All spectra shown in this work were corrected by inner filter effects using the following equation:^22, 23^

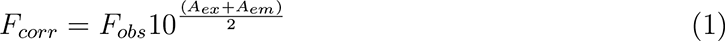

where *F_corr_*is the corrected fluorescence intensity, *F_obs_* is the experimentally measured intensity, *A_ex_* is the absorbance at the excitation wavelength (295nm) and *A_em_* is the absorbance at the emission.

A quantitative analysis of suppression is possible using the Stern-Volmer model fit. Some equations are used for the Stern-Volmer adjustment. We used the equation with the most satisfactory results, compatible with those obtained through other experimental techniques:^24, 25^

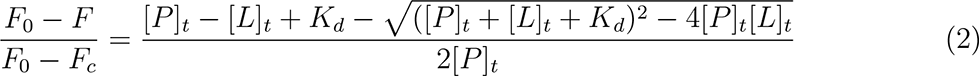

Where [*P*]*_t_* is the protein concentration, *F*_0_ is the protein fluorescence integral in the absence of ligand, *F* is the fluorescence integral in the presence of ligand, [*L*]*_t_* is the ligand concentration, *F_c_* is the integral of the remaining fluorescence. *K_d_* is the fit parameter, dissociation constant. The inverse of the dissociation constant is defined as the binding constant, *K_a_*=1/*K_d_*.

### Circular Dichroism

CD measures were performed with a JASCO-815 CD spectropolarimeter within a wavelength range of 195–260 nm. For recording the CD spectrum, a scan speed of 100 nm/min and a spectral bandwidth of 2.00 nm, were maintained as parameters. A quartz cuvette with a 10mm path length was used for the entire study. The following equation was used to convert the Theta machine units (*θ*, millidegrees) in Mean Residue Ellipticity (MRE, *degrees.cm*^2^*.dmol^−^*^1^*.residue^−^*^1^):

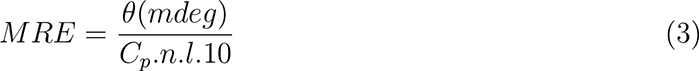

where C*_p_* = molar concentration of the protein, n = number of amino acid residues, and l = cell path length. The *α*-helical content was calculated using the following equation adjusted for the BSA protein:^26^

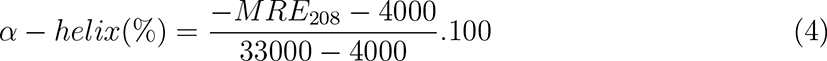

where *MRE*_208_ is the mean residue ellipticity at 208 nm.

### Small Angle X-ray Scattering

SAXS measurements were performed on the SAXS1 beamline of LNLS, using a Pilatus 300K detector positioned at *∼*1000mm from the sample. All measurements were performed using a sample holder with a path length of 1 mm and made of two mica windows. All SAXS measurements were performed at 22*±*1*^◦^*C. The SAXS intensity of an isotropic solution of monodisperse particles is given by:

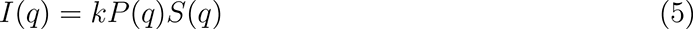

where k is a factor related to the instrumental effects and to the particle number density, *q* is the scattering vector, *P* (*q*) is the particle form factor, and *S*(*q*) is the structure factor. *S*(*q*) *∼* 1 for diluted (not interacting) systems, so *I*(*q*) is proportional to *P* (*q*). In the current work, we use the Guinier approximation to calculate the radius of gyration (*R_g_*) as follows:

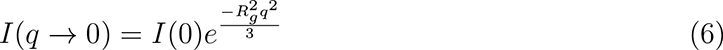

Noteworthy, Guinier’s law can only be applied in the *q_max_R_g_≤* 1.3 range (for approximately globular proteins) and for non-interacting systems.^27^

### Molecular Dynamics simulations

We have performed molecular dynamics (MD) simulations of BSA in aqueous 1-methyl-3-tetradecylimidazolium chloride ([C_14_MIM][Cl]) solutions in 4 different concentrations: 1 BSA and 0 [C_14_MIM][Cl] (control simulation), 1 BSA and 10 [C_14_MIM][Cl], 1 BSA and 100 [C_14_MIM][Cl], 1 BSA and 200 [C_14_MIM][Cl]. The simulated systems were composed of 1 BSA, 16 sodium counterions to neutralize the protein charge; and N water molecules in a very large cubic box with a side length around 16 nm (N = 117000 for 1 BSA/0 [C_14_MIM][Cl], N = 116697 for 1 BSA/10 [C_14_MIM][Cl], N = 115623 for 1 BSA/100 [C_14_MIM][Cl], N = 114243 for 1 BSA/ 200). The simulations were run in an NPT ensemble at 1 atm pressure and 298.15 K temperature. The velocity rescaling thermostat^28^ was used to regulate temperature with a coupling constant of 0.1 ps, and the Berendsen barostat^29^ was used to control pressure with a coupling constant of 2 ps, to create the NPT ensemble. All interactions were calculated within a 14-cutoff radius. With cubic interpolation, a 14-bit Fourier spacing, and the smooth particle-mesh Ewald approach, an extended correction for electrostatic interactions was taken into account. The leapfrog algorithm was used to integrate the equations of motion.^30^ The LINC**^?^** method was used with a time step of 2 fs and constraints in all H bonds. At each system level, the center of mass motion was linearly removed.The simulations were previously thermalized for 10 ns and the production phase consisted of 300 ns for the simulations containing 0 or 10 [C_14_MIM][Cl] and 500 ns for the simulations containing 100 or 200 [C_14_MIM][Cl]. The Gromos54A7^31, 32^ force field was adopted for BSA and [C_14_MIM][Cl] and the SPC/E^33^ model was adopted for water. The simulations were performed in Gromacs 5.1.4 package.^34^

## Results

### Fluorescence quenching and shift of BSA by ILs

The fluorescence spectra of BSA at 30µM in the absence and presence of increasing ILs concentrations are shown in Fig. 1A. Fig. 1A shows that the fluorescence of BSA gradually decreases (quenching) and shifts (blue-shift) with [C_14_MIM][Cl] addition. This behavior is similar to others reported in the literature with ionic and non-ionic surfactants. ^35, 36^ Fluorescence quenching requires proximity between the fluorophore and its quencher, ^37^ so we can suggest that there is an interaction between the ILs and BSA, at least with one of the tryptophans present in the protein at positions W134 and W213.

**Figure 1:**
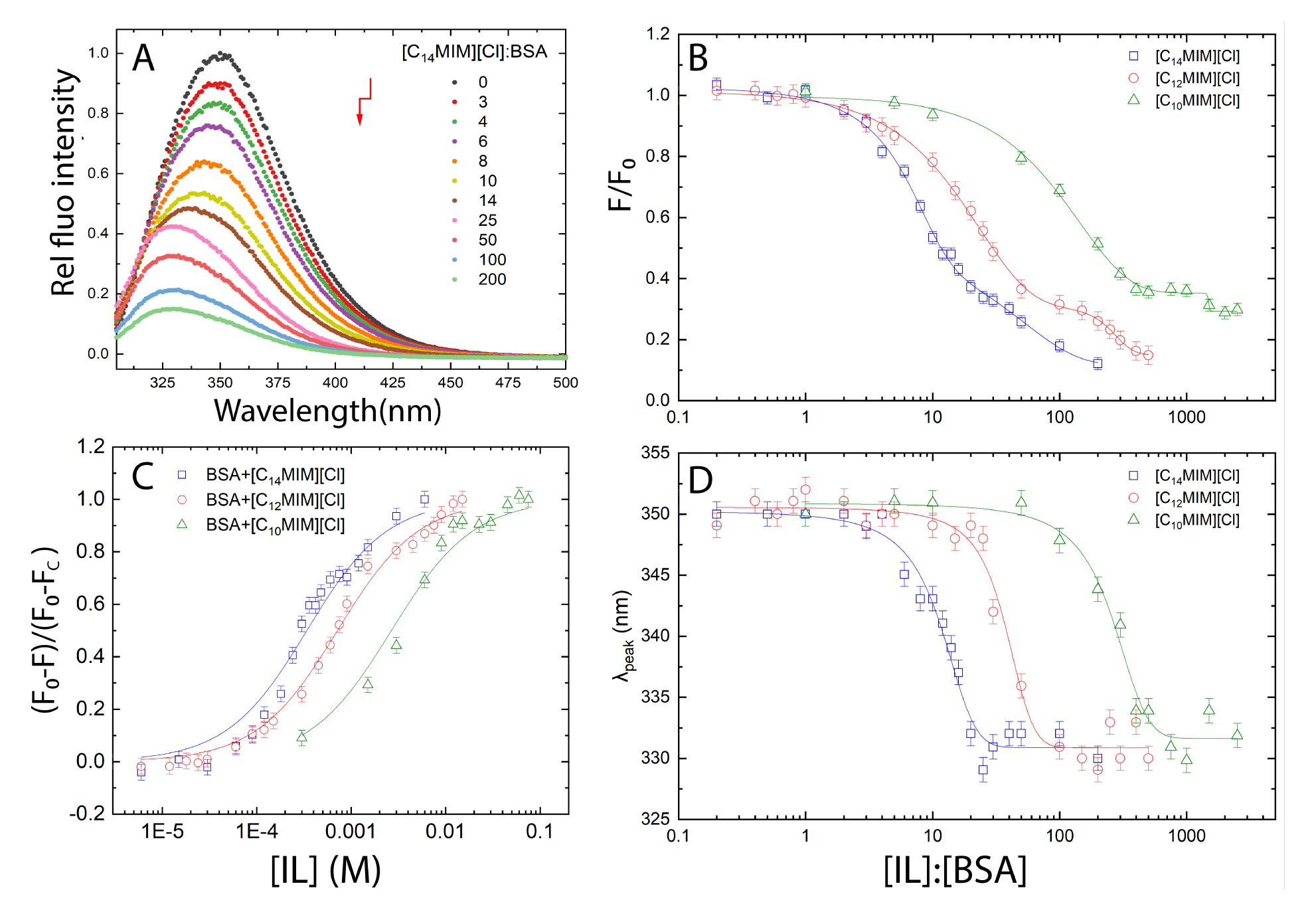
(A) Relative fluorescence emission spectra of 30 µM BSA (λ_*ex*_ = 295 nm) in the absence and presence of gradual addition of [C_14_MIM][Cl]. Molar ratios [IL]:[BSA] are in the legend (IL concentration increases along the arrow). (B) Ratio between the total fluorescence intensity (the integral under each fluorescence spectra) in the absence and presence of increasing IL concentration as a function of molar ratio [IL]:[BSA] on a linear-log scale. (C) Stern-Volmer plot as a function of IL concentration on a linear-log scale. (D) The wave-length at the maximum emission of fluorescence as a function of molar ratio [IL]:[BSA] on a linear-log scale.

The ratio between the total fluorescence intensity in the presence of IL and in the absence of IL (F/F_0_) is shown in Fig. 1B. F/F_0_ was calculated by dividing the integral of the fluorescence spectrum in the presence of ionic liquid by the integral of the fluorescence spectrum in the absence of ionic liquid. Stern-Volmer plot for the quenching of fluorescence intensity using Equation 2 is shown in Fig. 1C. Fig. 1D shows the wavelength of the maximum emission of fluorescence, as a function of IL concentration.

Concerning the quenching effect, from Fig. 1B, we observed total fluorescence intensity decreased by 50% at the molar ratio of 12±1, 30±3 and 202±3 of [C_14_MIM][Cl]:BSA, [C_12_MIM][Cl]:BSA and [C_10_MIM][Cl]:BSA, respectively. This result indicates that the length of the alkyl chain plays a relevant role in the IL/BSA interaction, in other words, the longer the alkyl chain, the stronger the binding. In addition, that effect could be related to the hydrophobic characteristic of such interaction. The Stern-Volmner plot was adjusted according to the Equation 2 to obtain *K_a_* values, Fig. 1C.

*K_a_* is larger as the carbon chain length increases as follows: (30 *±* 2).10*M ^−^*^1^, (14.3 *±* 0.7).10*M ^−^*^1^ and (3.8*±*0.4).10*M ^−^*^1^ in the presence of [C_14_MIM][Cl], [C_12_MIM][Cl] and [C_10_MIM][Cl], respectively. The *K_a_* values were calculated in order to compare the interaction of the BSA with the 3 ILs, however, for a more reliable obtainment of the binding constant, other techniques are more adequate, such as isothermal titration calorimetry. Nevertheless, this result reinforces the importance of the hydrophobic component for the LI/W interaction.

Fig. 1D can be divided into 3 different regions, as follows. In the first region, there is a plateau around 350.5±0.5nm that extends, in a molar ratio (i.e., [IL]:[BSA]), from 0:1 to 2:1, from 0:1 to 8:1, and from 0:1 to 50:1 of [C_14_MIM][Cl]:BSA, [C_12_MIM][Cl]:BSA and [C_10_MIM][Cl]:BSA, respectively. In this first stage, at small ILs concentrations, there is no significant change in the emission peak wavelength. In the second region, however, there is a significant decrease in wavelength (i.e. a blue-shift), starting from 350.5±0.5nm and reaching the minimum value of 331±1nm for all ionic liquids (at the reported concentrations), which was quite unexpected (Fig. 1D). This second stage shows a blue shift in lower IL concentration the higher the alkyl chain. In the third region, it was observed a new plateau around 331±1nm, starting at molar ratios of 35:1, 100:1 and 1000:1 for [C_14_MIM][Cl], [C_12_MIM][Cl] and [C_10_MIM][Cl], respectively. This third stage evidences a saturation-like behavior in the interaction of the protein and the amphiphilic molecules.

The fluorescence results we found are in agreement with other studies reported in the literature on the interaction of albumins with surfactants or ionic liquids surfactants. The interaction of the nonionic surfactant Hecameg with BSA shows the same behavior in three stages.^35^ The first stage is ascribed to an initial non-cooperative binding process, the second to a cooperative binding, and the third to a saturation binding region.

The gradual blue shift with increasing LI concentration may indicate a more hydrophobic environment near tryptophan.^37^ However, Vivian and Callis^38^ demonstrated that the proximity of charges or dipoles can alter tryptophan emission for both shorter and longer wavelengths, depending on the orientation. Therefore, a deeper discussion will be done along with the results of molecular dynamics.

### The Influence of IL on the protein tertiary structure

In order to get structural information on the IL/protein interaction, we performed SAXS measurements on 60µM of BSA in the absence and presence of increasing IL concentration. The respective scattering curves are presented in Fig. 2A. The curves are vertically shifted to emphasize the different scattering profiles, mainly in the high q range.

**Figure 2:**
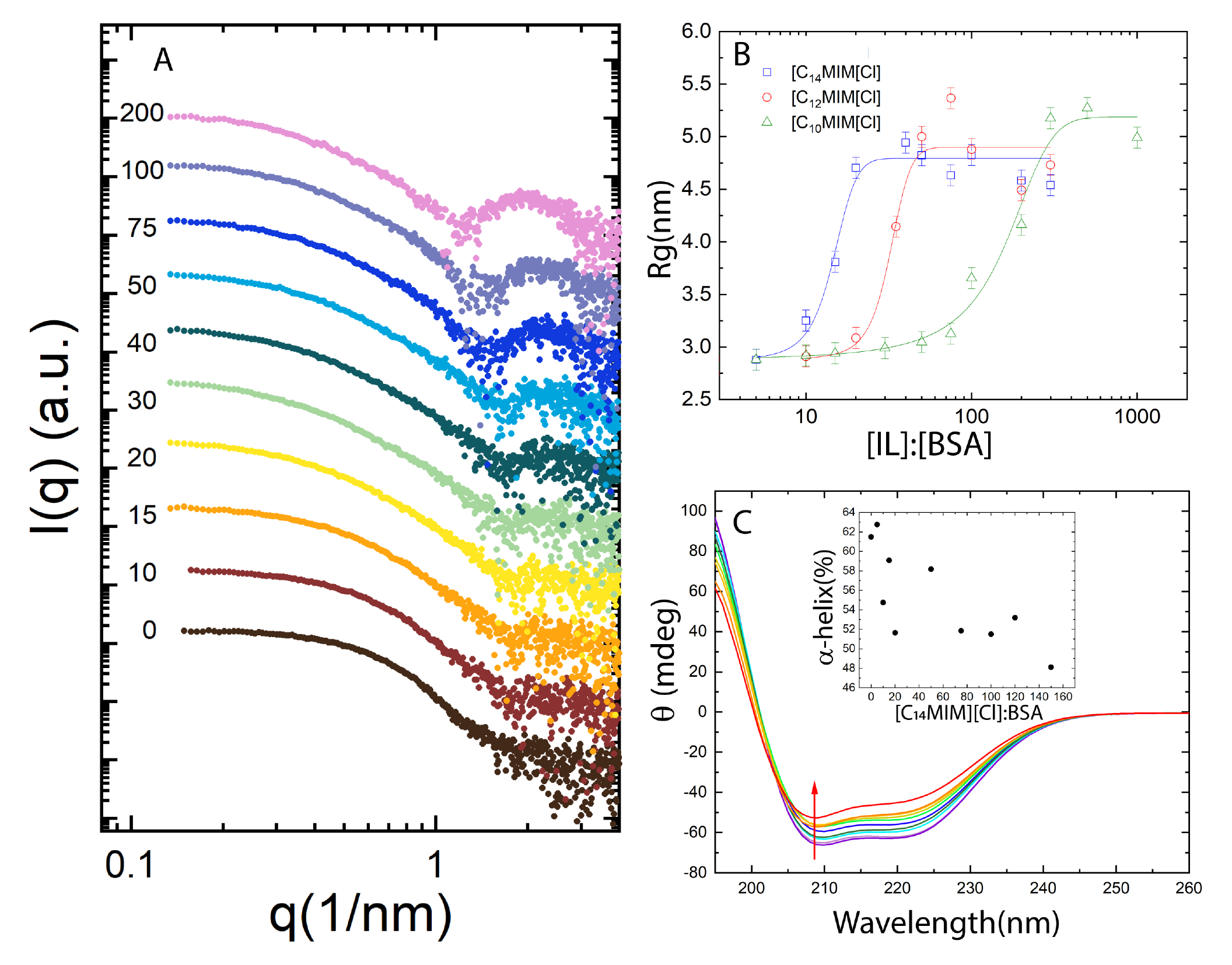
(A) SAXS data of 60 µM BSA in the absence and presence of gradual addition of [C_14_MIM][Cl]. Molar ratios [IL]:[BSA] are on the left. Curves are displaced vertically for clarity. (B) Radius of gyration in the absence and presence of increasing IL concentration as a function of molar ratio [IL]:[BSA] on a linear-log scale. The solid curve is used to guide the eyes. (C) Circular dichroism spectra of BSA in the absence and presence of gradual addition of [C_14_MIM][Cl]. Insert graph shows %*α − helix* content as a function of molar ratio [C_14_MIM][Cl]. IL concentration increases along the arrow.

Fig. 2A shows the formation of a peak at *q* = 0.23Å*^−^*^1^ as the molar ratio [C_14_MIM][Cl]:BSA increases. Such a peak is a fingerprint of micellar form factor from SAXS measurements. Then the data were analyzed through *R_g_* using the Guinier approximation and being shown in Fig. 2B.

Fig. 2B reveals *R_g_*increase from (29±1)Å in the absence of IL to (46±1)Å in the presence of 2.4mM of [C_14_MIM][Cl], and after such a concentration a plateau is reached. In the absence of IL, the *R_g_* value is in good agreement with the reported 30Å for the native crystallographic structure.^39^ This demonstrates that BSA is unfolding with IL interaction, saturating at 2.4mM of [C_14_mim][Cl]. This value is higher than the critical micellar concentration of this IL in buffer (1.0±0.1 mM calculated with superficial tension – data not shown), therefore further binding of the IL on the protein does not occur and there is coexistence of micelles and BSA-LI complex.

### ILs are able to slightly decrease the *α*-helix content of BSA

CD is an important tool to investigate modifications in proteins’ secondary structure. BSA is a protein that has mostly *α*-helix structures in its native state, which is reflected in two negative minima in the CD spectrum at 208nm and 222nm wavelengths.^40^ The literature reports values between 60% and 67% of *α*-helix for native BSA.^41^ The %*α*-helix is maximum when the BSA is in its native state and decreases in its unfolded state.^42^

The CD spectrum is shown in Fig. 2C. It is possible to observe that, in general, the minimum value at 222nm increases as [C_14_MIM][Cl] concentration increases (see red arrow on Fig. 2C), so there is an indication that the ionic liquid modifies the secondary structure even slightly. The inset in Fig. 2C shows the *α*-helix of BSA calculated using Equations 3 and 4. BSA in its native state, in the absence of IL, presents 61.5% of *α*-helix, which is in good accordance with the literature. Note that there is a downward trend in the *α*- helical content, reaching a 48.1% at 150 molar ratios. Such values indicate that the protein is unfolding. [C_12_MIM][Cl] and [C_10_MIM][Cl] have a similar, but smoother behavior. The lowest *α − helix* found with [C_12_MIM][Cl] and [C_10_MIM][Cl] are 52.3% and 54.6% at 300 and 50 molar ratios respectively.

Chakraborty et al.^19^ studied physicochemical and conformational changes in BSA-surfactant systems using different biophysical approaches, like Circular Dichroism, viscosimetry, turbid-ity, and Isothermal Tritation Calorimetry (ITC), and surface tension measurements. They reported two interesting physical constants, namely *c*_0_, and *c*_1_. The first one is the surfactant concentration where the system turbidity begins to increase, i.e., the system somehow is starting to change (surely, these are quite small numbers) due to the surfactant-protein interaction. The second one, on the other hand, is related to the surfactant concentration, where the surface tension reaches a plateau, i.e., all the added surfactants beyond this point are probably making micelles in the bulk solution, this should be understood as a saturation point. Interestingly, They reported a value of 67% of *α − helix* content for BSA in the absence of surfactant, in good accordance with the literature.^18^ On the other hand, they reported values of *α − helix* for BSA as 46%, 46% and 49% in the presence of the cationic hexadecyltrimethylammonium bromide (CTAB), and tetradecyltrimethylammonium bromide (TTAB), and Dodecyltrimethylammonium bromide (DTAB), respectively, at surfactant concentrations smaller than *c*_0_, i.e., at small cationic surfactant concentrations. Nevertheless, the authors also report values for *α − helix* for BSA of 12%, 16% and 15%, for CTAB, TTAB, and DTAB, respectively, beyond *c*_1_, i.e., at surfactant saturation conditions. Such a fact indicates that at higher concentrations these cationic surfactants can really unfold BSA, up to a certain point. Therefore, in terms of the denaturing potential of BSA, ionic liquids are comparable to CTAB, TTAB and DTAB (cationic surfactants) before *c*_0_, i.e., at small ionic liquid concentration. Unfortunately, we were not able to measure larger IL concentrations using CD, due to the chemical nature of our IL, which absorbs UV light.

### Investigating the location of IL on BSA surface using molecular dynamic simulations

To shed some light on the IL effects on BSA structure and electronic properties, we have performed MD simulations of BSA in pure aqueous solution and at 3 different concentrations of [C_14_MIM][Cl] ionic liquid (1 BSA/10 IL, 1 BSA/100 IL and 1 BSA/200 IL). The Root Mean Square Deviation (RMSD) of BSA, for the backbone or all atoms, did not show significant changes in the free IL simulation or in those ones at the 1/10 and 1/100 BSA/IL concentrations (Fig. S1). But, at the highest IL concentration, 1 BSA/200 IL, it has increased around 5% between 200 and 300 ns of simulation, equilibrating after that. Such structural changes are mainly related to the protein radius of gyration enhancement (Fig. 3), from 2.67*±*0.17 nm in free IL simulation to 2.89*±*0.03 nm in 1 BSA/200 IL simulation, that probably occurs in the region of residues 550 to 583, as revealed by the Root Mean Square Fluctuations (RMSF) analysis (Fig. S2). Although the calculated changes in the BSA radius of gyration are smaller than the experimental observations, in the absence and presence of C_14_MIM at 3mM has increased from 2.9 to 4.5 nm, respectively, the theoretical tendency is in agreement with our experimental findings, demonstrating that at high ionic liquid concentrations, the interaction between BSA and the IL can indeed cause an increase of the protein radius of gyration.

**Figure 3:**
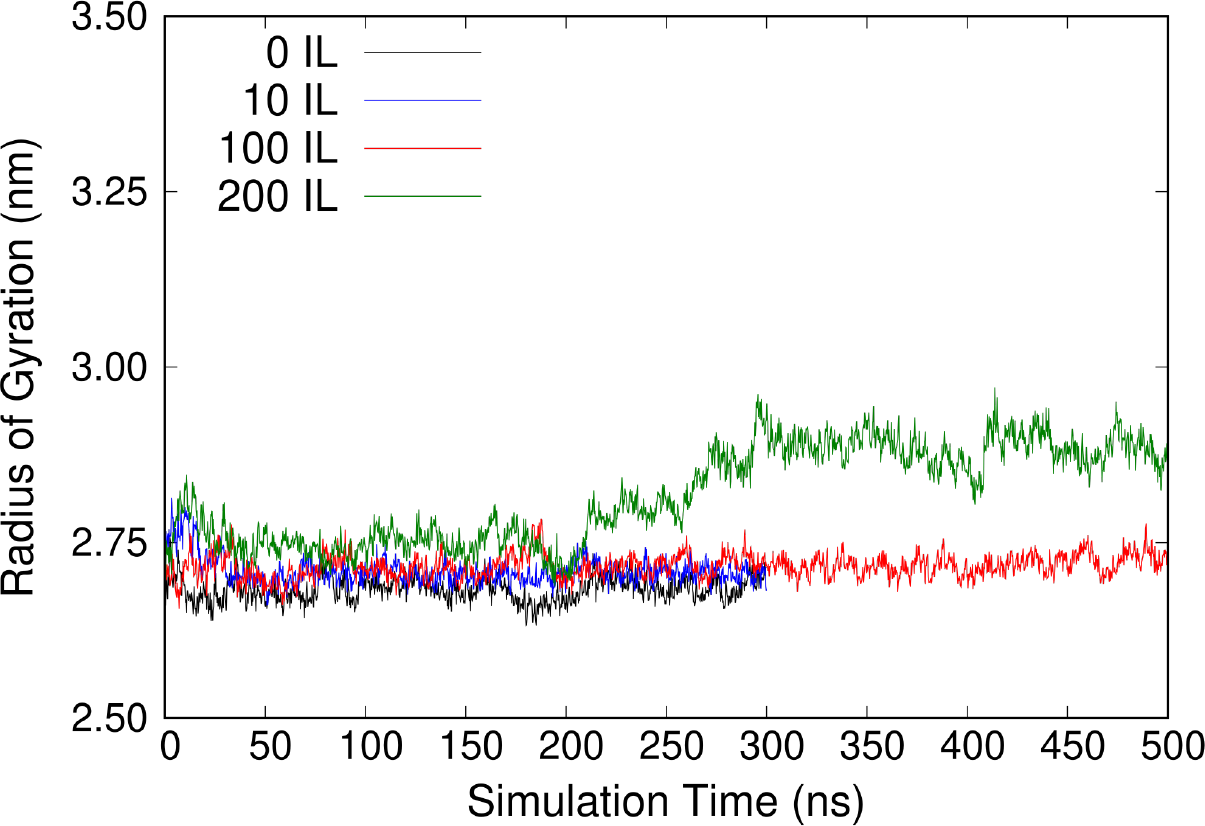
Radius of gyration of BSA in pure water (in black) and in IL/water solutions at different concentrations (1 BSA/10 IL in blue, 1 BSA/100 IL in red, 1 BSA/200 IL in green).

The solvation shells around BSA are quite different in the simulations at high (100 or 200 IL per BSA) and lower ionic liquid concentrations (0 or 10 IL per BSA). At the highest concentrations, the cationic C_14_MIM molecule causes strong dehydration of the protein, where lots of water/BSA hydrogen bonds are broken (Fig. 4b) and a considerable area around the protein is occupied by the cationic C_14_MIM molecules. After equilibration, the number of BSA/water hydrogen bonds is approximately 1192*±*23, 1160*±*23, 985*±*24 and 960*±*21 for the systems with 0, 10, 100, and 200 IL. It means that at the highest IL concentration around 250 BSA/water hydrogen bonds have disappeared during the simulation. The solvent distribution around the protein can also be mapped by the radial distribution functions between BSA and the ionic liquid and water molecules (Fig. S3). The integral of such functions shows that the number of C_14_MIM molecules surrounding the protein increases accordingly to the concentration of IL in the solution. For example, up to a distance of 2 nm between the centers of mass of the protein residues and the C_14_MIM molecules, it is found out a total of 9, 7 and 1 C_14_MIM molecules surrounding the protein at the concentrations 1 BSA/10 IL, 1 BSA/100 IL and 1 BSA/200 IL, respectively. These numbers increase to 26, 19, and 3 for a distance of 3 nm and to 70, 53, and 8 for a distance of 5 nm, respectively. Fig. 4c-d shows MD snapshots of the C_14_MIM distribution around BSA for the 1 BSA/200 IL simulation at 50 and 500 ns. Those C_14_MIM molecules which are not found close to the protein solvent accessible area form small micellar aggregates containing 2, 3, and up to 12 C_14_MIM units.

**Figure 4:**
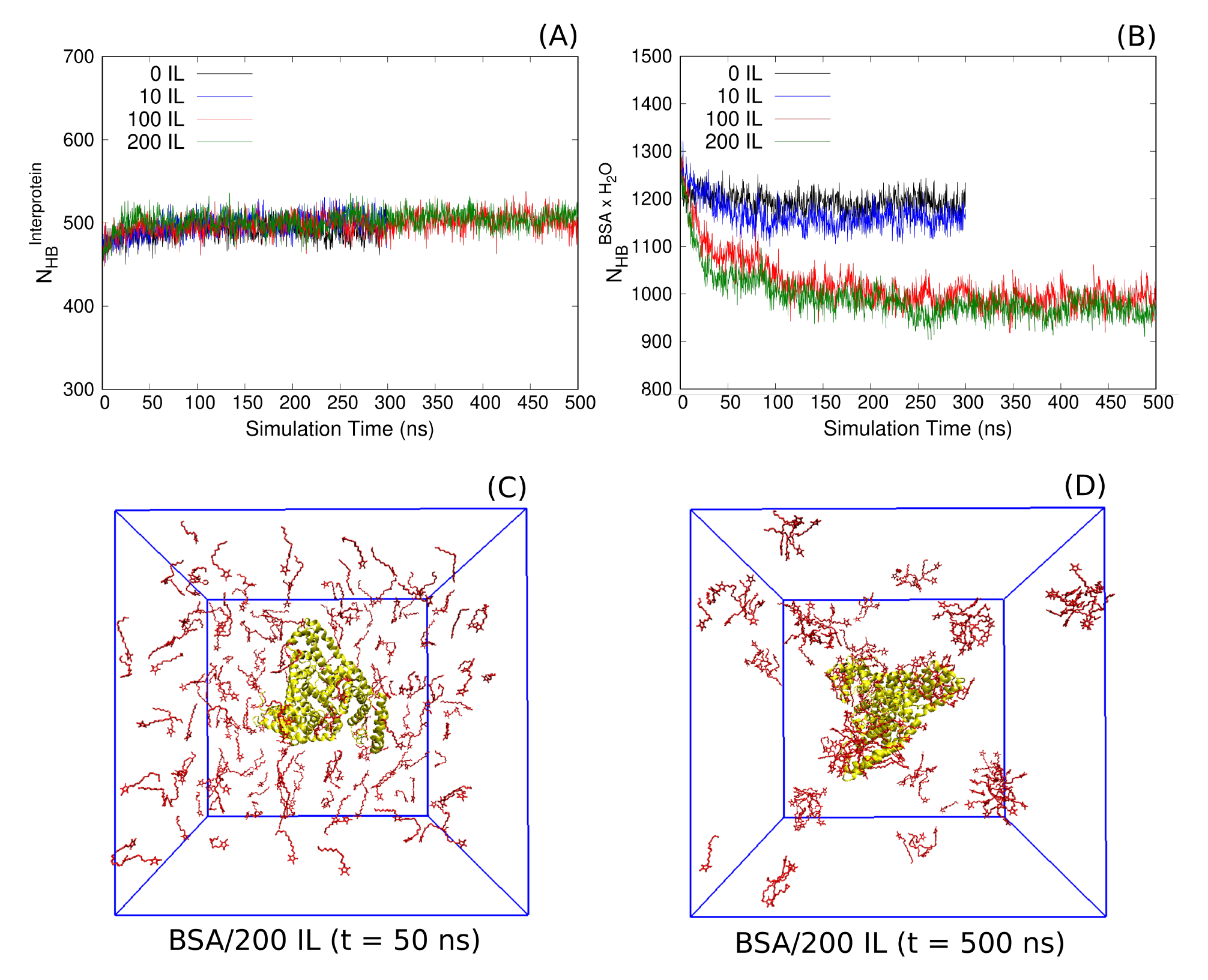
Number of intramolecular (A) and intermolecular (C) hydrogen bonds for simulations of BSA in pure water in black) and in ionic liquid (IL)/water solutions of different concentrations (1 BSA/10 IL in blue, 1 BSA/100 IL in red, 1 BSA/200 IL in green). (C) Snapshots extracted from the simulation of BSA/200IL showing the protein (in yellow) and the C_14_MIM molecules (in red), waters, and counter ions excluded in the images for better visualization.

On the other hand, the IL did not affect the secondary structure of BSA. The coil, bend, turn, alfa-helix, and 3-helix percentages were quite similar in any simulation (Fig. S4 and Tab. S1). The calculated percentage of *α*-helix was 67, 69, 68, and 69% in the systems with 0, 10, 100, and 200 IL, respectively. These numbers are close to our experimental prediction that was 66% and agrees with the theoretical observation that the number of BSA intermolecular hydrogen bonds does not change during simulations as well (Fig. 4a), being 501*±*13, 503*±*11, 504*±*11 and 505*±*11 in the systems with 0, 10, 100 and 200 IL.

Finally, we have analyzed the C_14_MIM distribution around the two Trp residues. Quite interestingly, in the simulations with 100 or 200 IL, a single C_14_MIM molecule was found close to the most exposed Trp of BSA (TRP1) interacting with it in a type of *π − π* stacking configuration, with the imidazole ring of C_14_MIM parallel to the TRP1 pyrrole and benzene rings, with a binding energy of −6.5*±*2.0 kcal/mol. Such configuration was observed before the first 10 ns (or 60 ns) of simulation for the system with 200 IL (or 100 IL) and remained during the entire simulation without C_14_MIM exchanges. The same was not observed for the buried Trp (TRP2), which was found reasonably close only to the extremity of any C_14_MIM tail that has entered the protein structure. Fig. 5a shows the minimum distance between the tryptophan residues (TRP1 and TRP2) and the C_14_MIM molecules over the simulations. For the last 200 ns of simulation, the TRP1-C_14_MIM minimum distance are on average 0.30*±*0.10 (or 0.34*±*0.10) for the 1BSA/200IL (or 1BSA/100IL) system and the TRP2-C_14_MIM minimum distance are on average 0.58*±*0.22 (or 0.52*±*0.26) for the 1BSA/200IL (or 1BSA/100IL) system.

**Figure 5:**
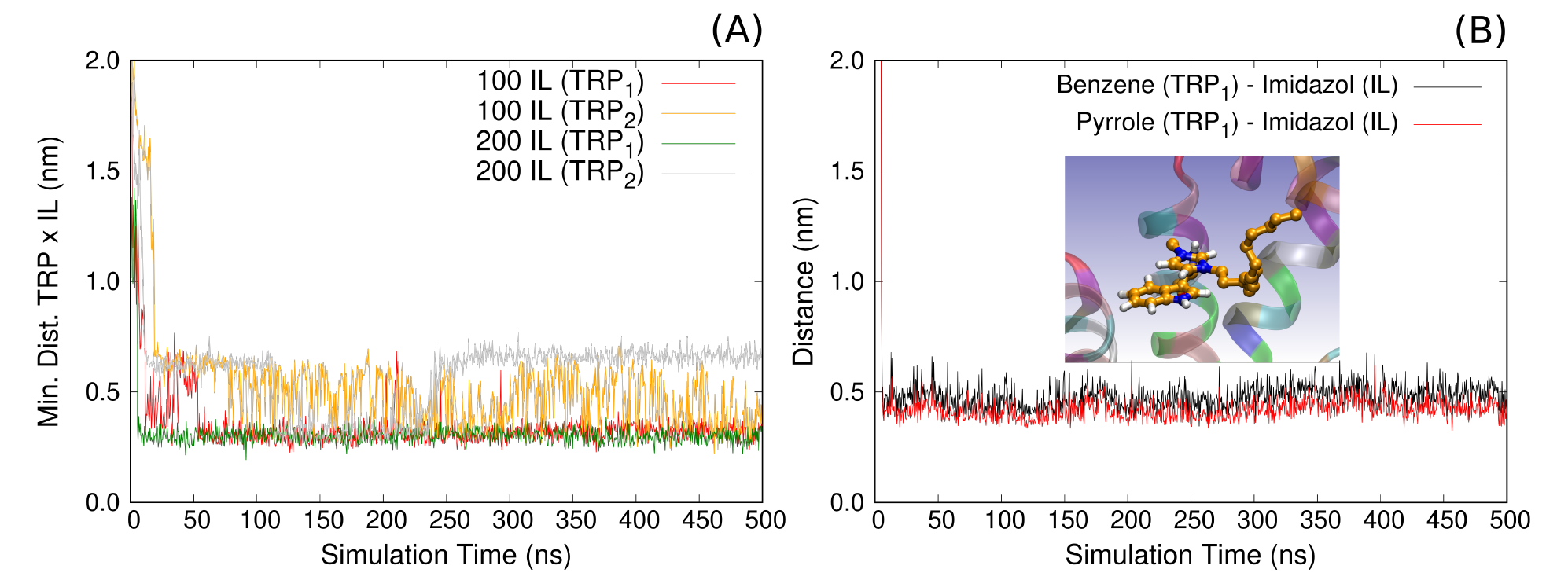
(A) Minimum distance between the tryptophan residues (TRP1 and TRP2) and the C14 IL molecules for BSA in ionic liquid/water solutions of different concentrations (1 BSA/100 IL and 1 BSA/200 IL). (B) Center of mass distances between the benzene and pyrrole rings of TRP1 and the imidazole ring of C14 for BSA in ionic liquid/water solution at 1 BSA/200 IL concentration.

The TRP1/C_14_MIM *π − π* stacking interaction was a surprising result. Analyzing the MD configurations, we observed that the center of mass distance from the pyrrole TRP1 ring to the imidazole C_14_MIM ring (0.41*±*0.2 nm) is approximately 0.9 Å smaller than the distance from the benzene TRP1 ring to the imidazole C_14_MIM ring (0.50*±*0.2 nm, see Fig. 5B). In other words, the positive +1 charge of the C_14_MIM imidazole is closer to the pyrrole ring than to the benzene ring of TRP1. This is an important observation that has an impact on the spectroscopy properties of BSA in ionic liquid solution. Vivian and Callis^38^ reported theoretical results where they have predicted the fluorescence wavelengths of 19 tryptophans in 16 proteins, and demonstrated that a positive charge over the tryptophan pyrrole ring induces a blue shift on the emission spectra of the protein. Therefore, our MD simulations findings can explain the measured blue shift in the emission spectra of BSA at high IL concentrations.

We can suppose that IL, at least in high concentrations, induces a structural change in BSA, increasing its radius of gyration. The regions within the BSA structure that most contribute to such modifications can be identified from the RMSF as a function of the position of the BSA residue, as shown in Fig. S4.

Through RMSF analysis of the BSA structure, we can notice that there are more flexible and dynamic regions within the structure than others, even in the absence of LI. We would like to highlight the 110-120 region for BSA in the presence of 100 molecules of LI and the 500-535 and 560-583 regions for BSA in the presence of 200 molecules of LI.

In the RMSF plot, we can observe that the main peaks are in the *α*-helices of subdomain III. Subdomain III, in addition to possessing important binding sites, is also the subdomain that undergoes the most modifications during the protein denaturation process due to changes in pH^43^.^44^

## Discussions

These results suggest that the interaction of BSA with ILs can be divided into three stages, as shown in Fig. 6. The first stage is characterized by native protein, when electrostatic forces were predominating, provided by the opposing charges on the surface of the proteins and the polar head of the ILs. The approximation of the IL to the surface of the protein allows the interaction between the hydrophobic sites of the protein with the carbon chains of the ILs. The hydrophobic effect is a relevant contributor to the interaction in the second stage, potentiating the formation of the protein-IL complex. In the last stage, the interaction between protein and IL decreases. The formation of ILs micelles in solution starts and occur the binding saturation of IL to proteins.

**Figure 6:**
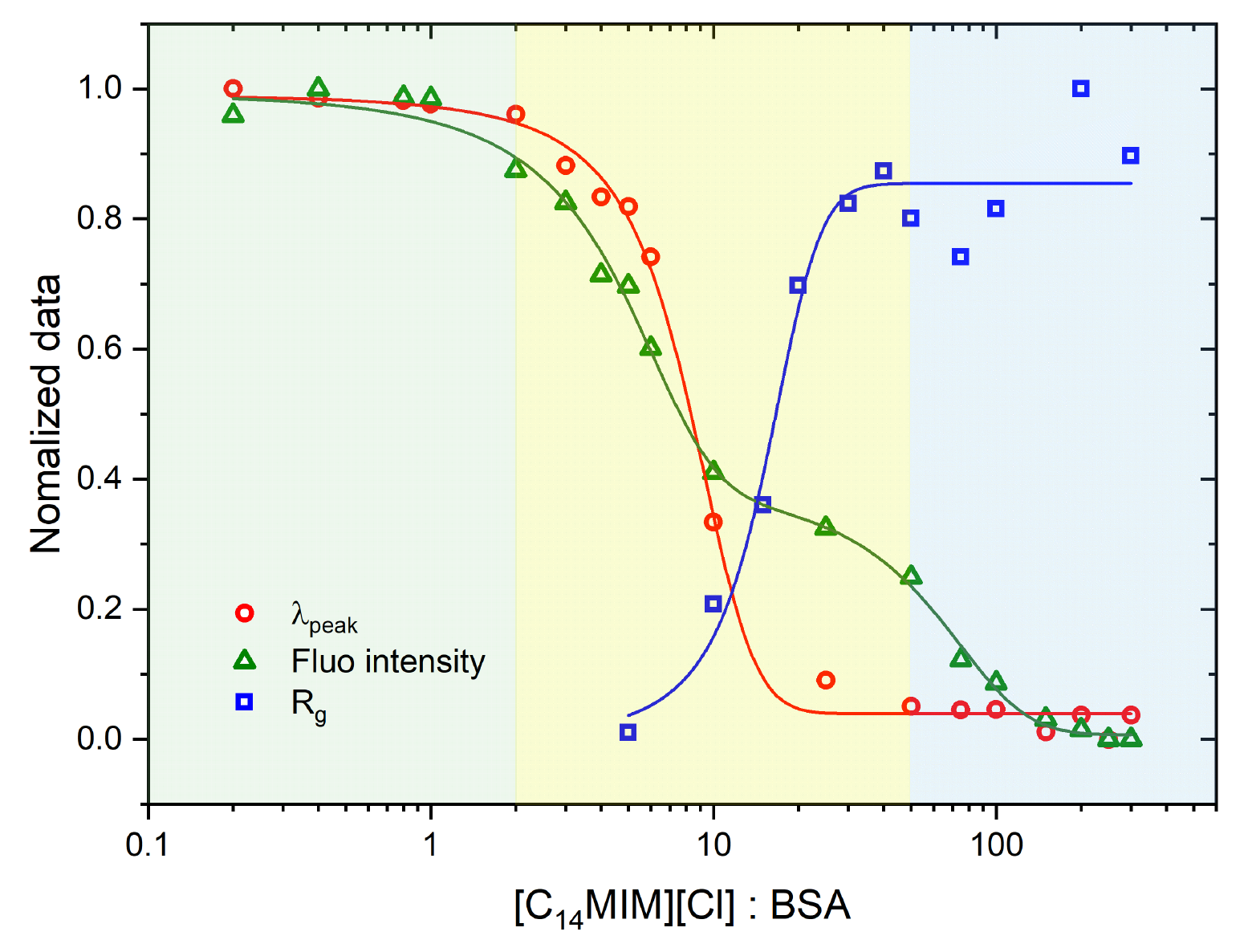
Normalized data from 0 to 1 of fluorescence shift, quenching and radius of gyration

The results of fluorescence spectroscopy obtained for BSA in the presence of [C_14_MIM][Cl], [C_12_MIM][Cl] and [C_10_MIM][Cl] show an interaction between the protein and the IL, more specifically near the tryptophan’s protein, inducing a quenching and a blue-shift. Analysis of radius of gyration from SAXS curves shows BSA unfolding with IL interaction. These effects indicate that the interaction of BSA with IL is divided into three stages. The interaction in the first stage is predominantly due to the electrostatic forces provided by the opposing charges on the surface of the proteins and the polar head of the ILs, in this case, the proteins are still in their native form. In the second stage, higher IL concentrations induce the unfolding of the proteins and therefore. The hydrophobic effect is a relevant contributor to the interaction. The accessibility of hydrophobic sites, by unfolding, potentiates the formation of the complex protein IL. Finally, in the third stage, IL micelles start to form, and, therefore, interaction with protein reaches a saturation point.

In their study, Gurbir Singh and Tejwant Singh Kang^45^ looked at how the BSA protein self-assembled in aqueous solutions of functionalized and unfunctionalized ionic liquids. Depending on the structure and concentration of the ILs, the effects on the self-assembled BSA generated by ce[C12MIM][Cl], ce[C12AMIM][Cl], and ce[C12EMIM][Cl] are all different. The authors saw large, chaotic self-assembled BSA structures, ordered self-assembled structures like lengthy rods, and right-handedly twisted helical amyloid fibers. However, a distinct IL action mechanism in BSA was noted in the current work.

It is well known that disulfide bonds are chemically stable in the absence of a reduction agent, which is our case since the bond cannot be reduced by the ionic liquid. Another way to avoid a disulfite bond is to perform a mutation in the Cys residue, which was not performed in the present study. Thus, from our perspective, and under the experimental conditions used herein, it is straightforward to conclude that all the 17 disulfide bonds remain unaltered during the ionic liquid/protein interaction. Such a fact can justify the secondary structure of BSA even at higher ILs concentrations.

According to the data presented herein, it is possible to propose a mechanism of action for the studied imidazolium ionic liquids with BSA. On this ground, Fig. 7 shows a schematic representation of the primary and secondary structure of BSA. Such a figure was assembled on the PDB website using the 4F5S entry for the protein. Figure 7 shows in the first line the protein residue number (ranging from 1 - 583), as well as, the regions with *α−helix* structures (the red rectangles in the second line of the figure) and most interestingly, the connection between each Cysteine pair for the 34 BSA cysteine residues. Moreover, in the last line, we have the position of Domain I (from residue 1 - 185), Domain II (from residue 186 - 378), and Domain III (from residue 379 −583). The data presented herein supports the model that the unfolding process starts with the increasing in the distance between Domains I and III, in such a way that the protein keeps its secondary structure. Moreover, it should be stressed that there are not cysteine bonds between domains. BSA has a heart-shaped structure, thus Domains I and III are almost in contact. The significance of this research relies in its potential applications in the pharmaceutical industry, where the use of ionic liquids as solvents, stabilizers, and drug delivery systems is gaining importance. Understanding the interactions between serum albumin and ionic liquids can provide valuable information for the development of more effective drug formulations and delivery systems with improved bioavailability and enhanced therapeutic outcomes.

**Figure 7:**
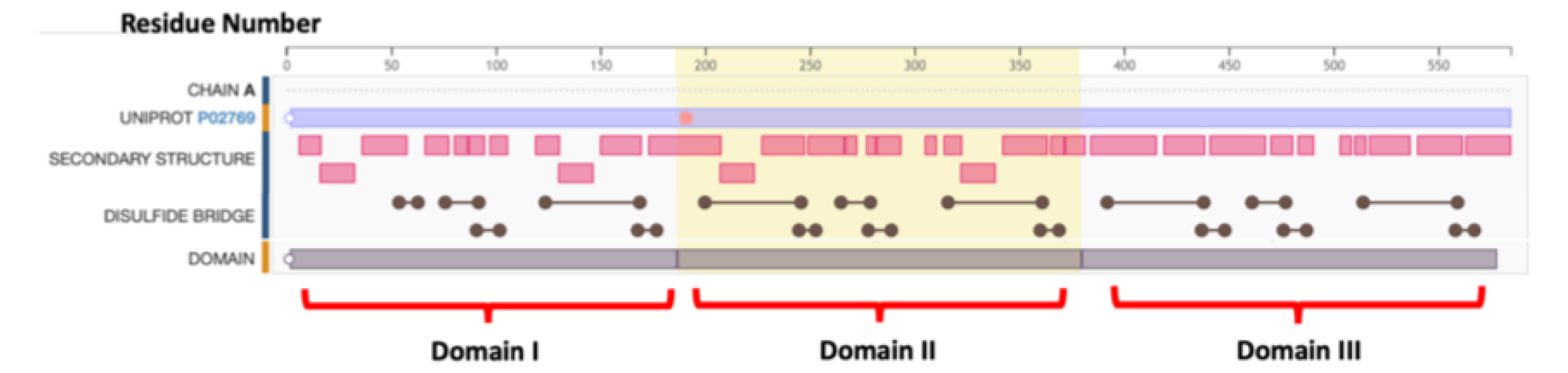
Squematic representation of Bovine Serum Albumin primary and secondary struc-ture, indicating the main structural features, like the Domain I (from residue 1 - 185), Domain II (from residue 186 - 378) and domain III (from residue 379 −583). It is also in-dicated in the figure the regions with *α − helix* structures (red rectangular boxes) and all the 17 disulfide bonds, involving the 34 cysteines (round circles connected with solid lines). These data represent the PDB entry 4F5S. This figure was created at www.pdb.org

## Acknowledgement

This work was supported by FAPESP (2015/15822-1, 2016/05019-0) and CNPq. Leandro R. S. Barbosa has research fellowships from CNPq (309418/2021-6). The authors thank the National Council for Scientific and Technological Development (CNPq) and Coordination for the Improvement of Higher Education (CAPES) for a research fellowship. This research used facilities of the Brazilian Synchrotron Light Laboratory (LNLS), part of the Brazilian Center for Research in Energy and Materials (CNPEM), a private non-profit organization under the supervision of the Brazilian Ministry of Science, Technology, and Innovations (MCTI). The SAXS1 beamline staff is acknowledged for their assistance during the experiments. We conducted the computational modeling using the High-Performance Computing (HPC) Cluster provided by the University of São Paulo.

## Supporting Information Available

This will usually read something like: “Experimental procedures and characterization data for all new compounds. The class will automatically add a sentence pointing to the infor-mation online:

## References

(1) Buettner, C. S.; Cognigni, A.; Schröder, C.; Bica-Schröder, K. Surface-active ionic liquids: A review. Journal of Molecular Liquids 2022, 347, 118160.

(2) Welton, T. Room-temperature ionic liquids. Solvents for synthesis and catalysis. Chemical reviews 1999, 99, 2071–2084.

(3) Mu, T.; Han, B. In Structures and Interactions of Ionic Liquids; Zhang, S., Wang, J., Lu, X., Zhou, Q., Eds.; Springer Berlin Heidelberg: Berlin, Heidelberg, 2014; pp 107–139.

(4) Greaves, T. L.; Drummond, C. J. Protic Ionic Liquids: Properties and Applications. Chemical Reviews 2008, 108, 206–237.

(5) He, P.; Liu, H.; Li, Z.; Liu, Y.; Xu, X.; Li, J. Electrochemical Deposition of Silver in Room-Temperature Ionic Liquids and Its Surface-Enhanced Raman Scattering Effect. Langmuir 2004, 20, 10260–10267.

(6) Kim, T.; Lee, H.; Stoller, M.; Dreyer, D.; Bielawski, C.; Ruoff, R.; Suh, K. High-performance supercapacitors based on poly(ionic liquid)-modified graphene electrodes. ACS Nano 2011, 5, 436–442.

(7) Attri, P.; Choi, E. H. Influence of Reactive Oxygen Species on the Enzyme Stability and Activity in the Presence of Ionic Liquids. PLOS ONE 2013, 8, 1–11.

(8) Park, S.; Viklund, F.; Huit, K.; Kazlauskas, R. Ionic Liquids as Green Solvent - Progress and Prospects; ACS Symposium Series; 2003; Vol. 856; pp 225–238.

(9) Pedersen, J. N.; Lyngsø, J.; Zinn, T.; Otzen, D. E.; Pedersen, J. S. A complete picture of protein unfolding and refolding in surfactants. Chemical science 2020, 11, 699–712.

(10) Otzen, D.; Pedersen, J. N.; Rasmussen, H. Ø.; Pedersen, J. S. How do surfactants unfold and refold proteins? Advances in Colloid and Interface Science 2022, 102754.

(11) Blanco, E.; Ruso, J.; Prieto, G.; Sarmiento, F. On relationships between surfactant type and globular proteins interactions in solution. Journal of Colloid and Interface Science 2007, 316, 37–42, cited By 20.

(12) Gull, N.; Sen, P.; Khan, R.; Kabir-Ud-Din, Spectroscopic studies on the comparative interaction of cationic single-chain and gemini surfactants with human serum albumin. Journal of Biochemistry 2009, 145, 67–77, cited By 37.

(13) Kaspersen, J. D.; Søndergaard, A.; Madsen, D. J.; Otzen, D. E.; Pedersen, J. S. Re-folding of SDS-Unfolded Proteins by Nonionic Surfactants. Biophysical Journal 2017, 112, 1609 – 1620.

(14) Otzen, D. Protein–surfactant interactions: A tale of many states. Biochimica et Biophysica Acta (BBA) - Proteins and Proteomics 2011, 1814, 562 – 591.

(15) Mackie, A.; Wilde, P. The role of interactions in defining the structure of mixed pro-tein–surfactant interfaces. Advances in Colloid and Interface Science 2005, 117, 3–13, A Collection of Papers from the International Workshop on Bubble and Drop Interfaces, Genoa, Italy, 25-28 April, 2004.

(16) Peters, T. All About Albumin; Academic Press: San Diego, 1995.

(17) Barbosa, L.; Ortore, M.; Spinozzi, F.; Mariani, P.; Bernstorff, S.; Itri, R. The importance of protein-protein interactions on the pH-induced conformational changes of bovine serum albumin: A small-angle x-ray scattering study. Biophysical Journal 2010, 98, 147–157, cited By 95.

(18) Itri, R.; Caetano, W.; Barbosa, L. R.; Baptista, M. S. Effect of urea on bovine serum albumin in aqueous and reverse micelle environments investigated by small angle X-ray scattering, fluorescence and circular dichroism. Brazilian journal of physics 2004, 34, 58–63.

(19) Chakraborty, T.; Chakraborty, I.; Moulik, S. P.; Ghosh, S. Physicochemical and Conformational Studies on BSA Surfactant Interaction in Aqueous Medium. Langmuir 2009, 25, 3062–3074, PMID: 19191571.

(20) Egorova, K. S.; Gordeev, E. G.; Ananikov, V. P. Biological Activity of Ionic Liquids and Their Application in Pharmaceutics and Medicine. Chemical Reviews 2017, 117, 7132–7189, PMID: 28125212.

(21) Han, Q.; El Mohamad, M.; Brown, S.; Zhai, J.; Rosado, C.; Shen, Y.; Blanch, E. W.; Drummond, C. J.; Greaves, T. L. Small angle X-ray scattering investigation of ionic liquid effect on the aggregation behavior of globular proteins. Journal of Colloid and Interface Science 2023,

(22) Sur, S. S.; Rabbani, L. D.; Libman, L.; Breslow, E. Fluorescence studies of native and modified neurophysins. Effects of peptides and pH. Biochemistry 1979, 18, 1026–1036, PMID: 34422.

(23) Mendoņca, A. F.; Rocha, A. F. C.; Duarte, A. C.; Santos, E. B. H. The inner filter effects and their correction in fluorescence spectra of salt marsh humic matter. Analytica chimica acta 2013, 788, 99–107.

(24) Bakar, K. A.; Feroz, S. R. A critical view on the analysis of fluorescence quenching data for determining ligand–protein binding affinity. Spectrochimica Acta Part A: Molecular and Biomolecular Spectroscopy 2019, 223, 117337.

(25) van de Weert, M.; Stella, L. Fluorescence quenching and ligand binding: A critical discussion of a popular methodology. J.Mol.Struct. 2011, 998, 144–150.

(26) Bhogale, A.; Patel, N.; Mariam, J.; Dongre, P.; Miotello, A.; Kothari, D. Comprehensive studies on the interaction of copper nanoparticles with bovine serum albumin using various spectroscopies. Colloids and Surfaces B: Biointerfaces 2014, 113, 276–284.

(27) Svergun, D. I.; Koch, M. H. Small-angle scattering studies of biological macromolecules in solution. Reports on Progress in Physics 2003, 66, 1735.

(28) Bussi, G.; Donadio, D.; Parrinello, M. Canonical sampling through velocity rescaling. The Journal of chemical physics 2007, 126, 014101.

(29) Berendsen, H. J.; Postma, J. v.; van Gunsteren, W. F.; DiNola, A.; Haak, J. R. Molecular dynamics with coupling to an external bath. The Journal of chemical physics 1984, 81, 3684–3690.

(30) Hockney, R. W.; Goel, S.; Eastwood, J. W. Quiet high-resolution computer models of a plasma. Journal of Computational Physics 1974, 14, 148 – 158.

(31) Huang, W.; Lin, Z.; van Gunsteren, W. F. Validation of the GROMOS 54A7 force field with respect to *β*-peptide folding. Journal of chemical theory and computation 2011, 7, 1237–1243.

(32) Schmid, N.; Eichenberger, A.; Choutko, A.; Riniker, S.; Winger, M.; Mark, A.; Van Gunsteren, W. Eur. Biophys. J. 2011.

(33) Berendsen, H.; Grigera, J.; Straatsma, T. The missing term in effective pair potentials. Journal of Physical Chemistry 1987, 91, 6269–6271.

(34) Van Der Spoel, D.; Lindahl, E.; Hess, B.; Groenhof, G.; Mark, A. E.; Berendsen, H. J. GROMACS: fast, flexible, and free. Journal of computational chemistry 2005, 26, 1701–1718.

(35) Hierrezuelo, J.; Nieto-Ortega, B.; Carnero Ruiz, C. Assessing the interaction of Hecameg® with Bovine Serum Albumin and its effect on protein conformation: A spectroscopic study. Journal of Luminescence 2014, 147, 15–22.

(36) Gelamo, E.; Silva, C.; Imasato, H.; Tabak, M. Interaction of bovine (BSA) and human (HSA) serum albumins with ionic surfactants: spectroscopy and modelling. Biochimica et Biophysica Acta 2002, 1594, 84 – 99.

(37) Lakowicz, J. R. Principles of Fluorescence Spectroscopy; Springer US, 2009.

(38) Vivian, J. T.; Callis, P. R. Mechanisms of Tryptophan Fluorescence Shifts in Proteins. Biophysical Journal 2001, 80, 2093–2109.

(39) Santos, S. F. A.; Zanette, D.; Fischer, H.; Itri, R. A systematic study of bovine serum albumin (BSA) and sodium dodecyl sulfate (SDS) interactions by surface tension and small angle X-ray scattering. Journal of colloid and interface science 2003, 262 *2*, 400–8.

(40) Corbin, J.; Méthot, N.; Wang, H. H.; Baenziger, J. E.; Blanton, M. P. Secondary structure analysis of individual transmembrane segments of the nicotinic acetylcholine receptor by circular dichroism and Fourier transform infrared spectroscopy. Journal of Biological Chemistry 1998, 273, 771–777.

(41) Halder, S.; Aggrawal, R.; Aswal, V. K.; Ray, D.; Saha, S. K. Study of refolding of a denatured protein and microenvironment probed through FRET to a twisted intramolecular charge transfer fluorescent biosensor molecule. Journal of Molecular Liquids 2021, 322, 114532.

(42) Halder, S.; Aggrawal, R.; Jana, S.; Saha, S. K. Binding interactions of cationic gemini surfactants with gold nanoparticles-conjugated bovine serum albumin: A FRET/NSET, spectroscopic, and docking study. Journal of Photochemistry and Photobiology B: Biology 2021, 225, 112351.

(43) El Kadi, N.; Taulier, N.; Le Huérou, J.; Gindre, M.; Urbach, W.; Nwigwe, I.; Kahn, P. C.; Waks, M. Unfolding and refolding of bovine serum albumin at acid pH: ultrasound and structural studies. Biophysical journal 2006, 91, 3397–3404.

(44) Baler, K.; Martin, O. A.; Carignano, M. A.; Ameer, G. A.; Vila, J. A.; Szleifer, I. Electrostatic Unfolding and Interactions of Albumin Driven by pH Changes: A Molecular Dynamics Study. The Journal of Physical Chemistry B 2014, 118, 921–930, PMID: 24393011.

(45) Singh, G.; Kang, T. S. Ionic Liquid Surfactant Mediated Structural Transitions and Self-Assembly of Bovine Serum Albumin in Aqueous Media: Effect of Functionalization of Ionic Liquid Surfactants. The Journal of Physical Chemistry B 2015, 119, 10573– 10585, PMID: 26230661.

