## Supplemental File for "Unveiling the three-step model for the interaction of imidazolium-based ionic liquids on albumin"

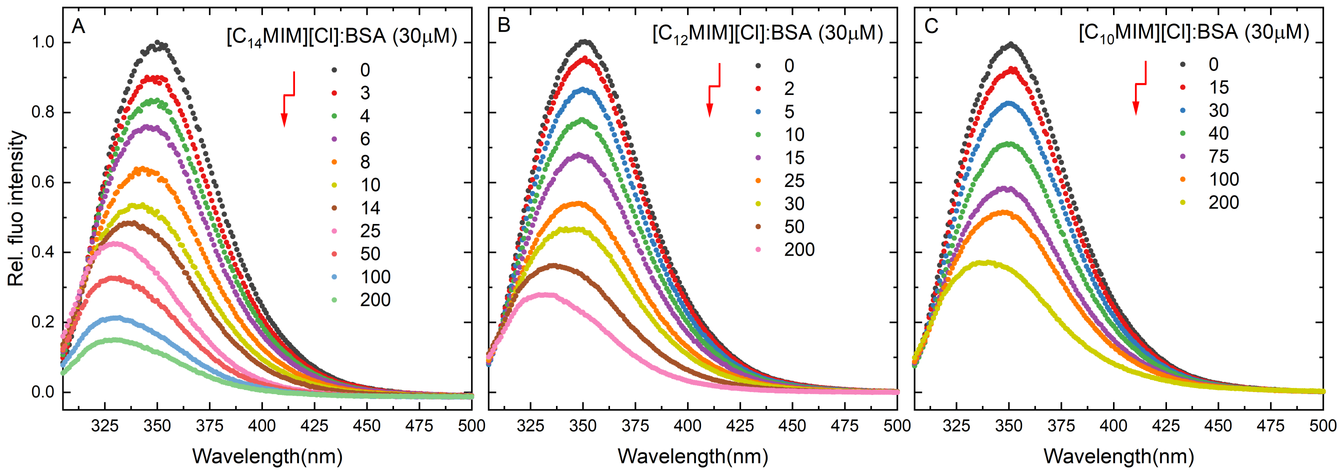


Figure 1: Relative fluorescence emission spectra of 30 $\mu$M BSA ($\lambda_{ex}$ = 295 nm) in the absence and presence of gradual addition of (A) [$C_{14}$MIM][Cl], (B) [$C_{12}$MIM][Cl], and (C) [$C_{10}$MIM][Cl]. Molar ratios [IL]:[BSA] are in the legend (IL concentration increases along the arrow).

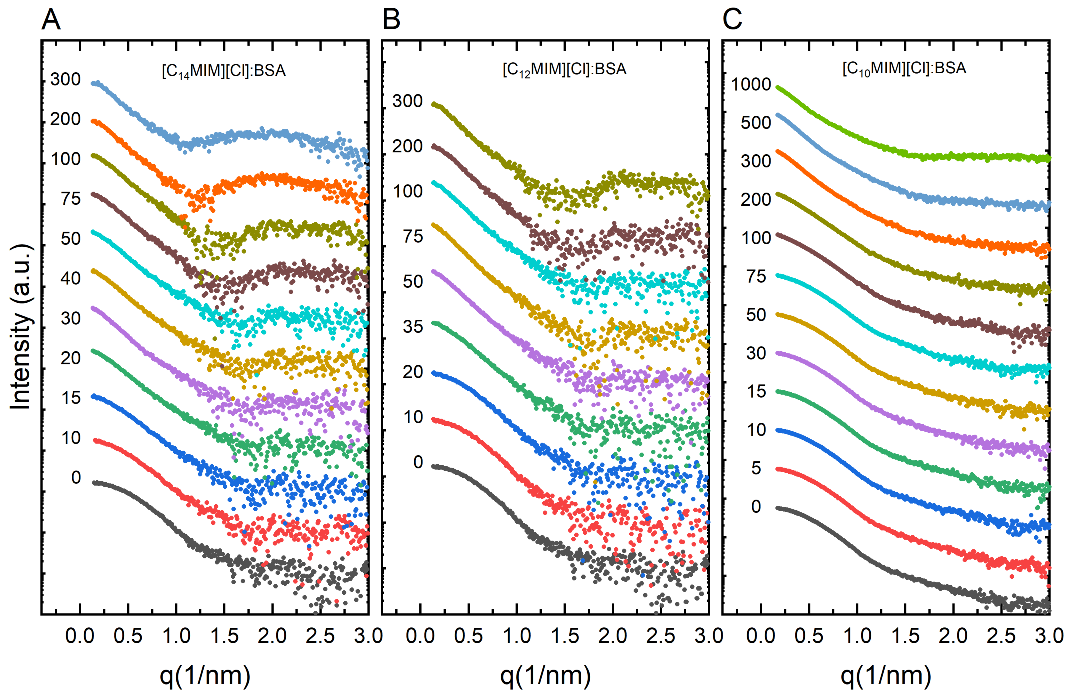


Figure 2: SAXS intensity of 60 $\mu$M BSA in the absence and presence of gradual addition of (A) [$C_{14}$MIM][Cl], (B) [$C_{12}$MIM][Cl], and (C) [$C_{10}$MIM][Cl]. Molar ratios [IL]:[BSA] are in the legend.

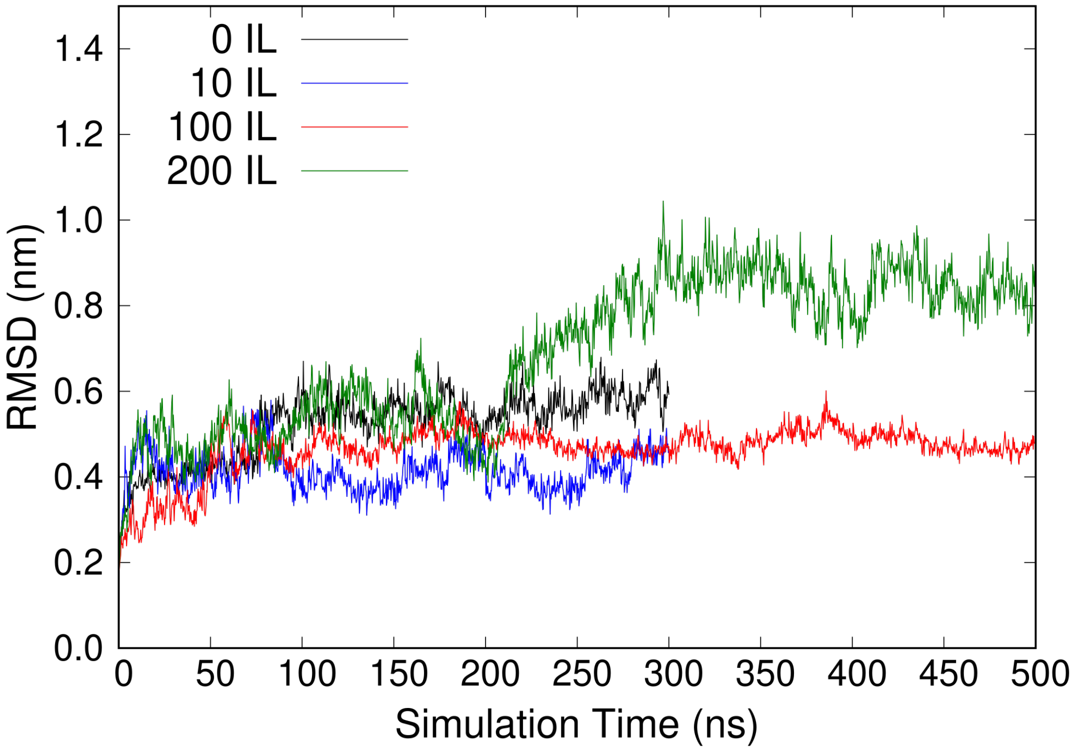


Figure 3: Root Mean Square Deviation (RMSD) of BSA backbone in pure water and in ionic liquid (IL)/water solutions of different concentrations (1 BSA/10 IL in blue, 1 BSA/100 IL in red, 1 BSA/200 IL in green). RMSD computed over the entire simulations on the reference of the first MD configuration.

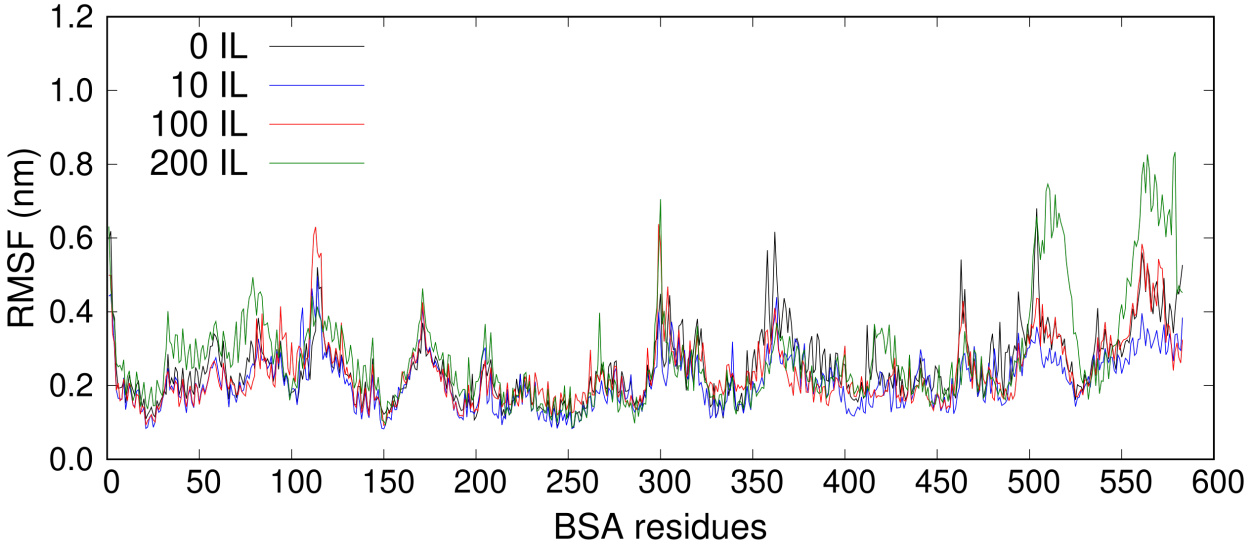


Figure 4: Root Mean Square Fluctuations (RMSF) of BSA residues in pure water and in ionic liquid (IL)/water solutions of different concentrations (1 BSA/10 IL in blue, 1 BSA/100 IL in red, 1 BSA/200 IL in green). RMSF computed over the entire simulations.

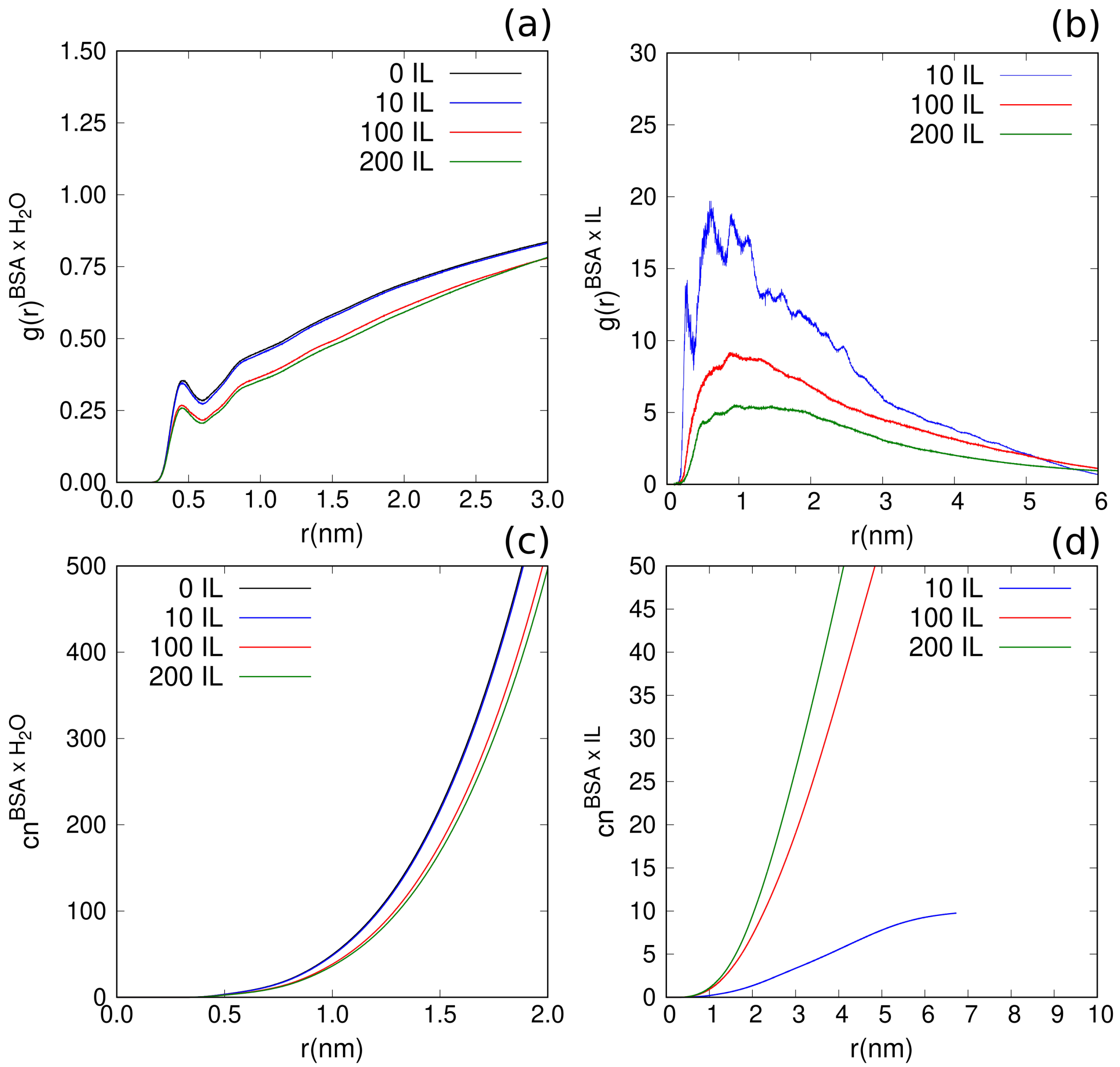


Figure 5: Radial distribution functions (g(r)) and coordination numbers (cn) between BSA and the ionic liquid/water solutions.

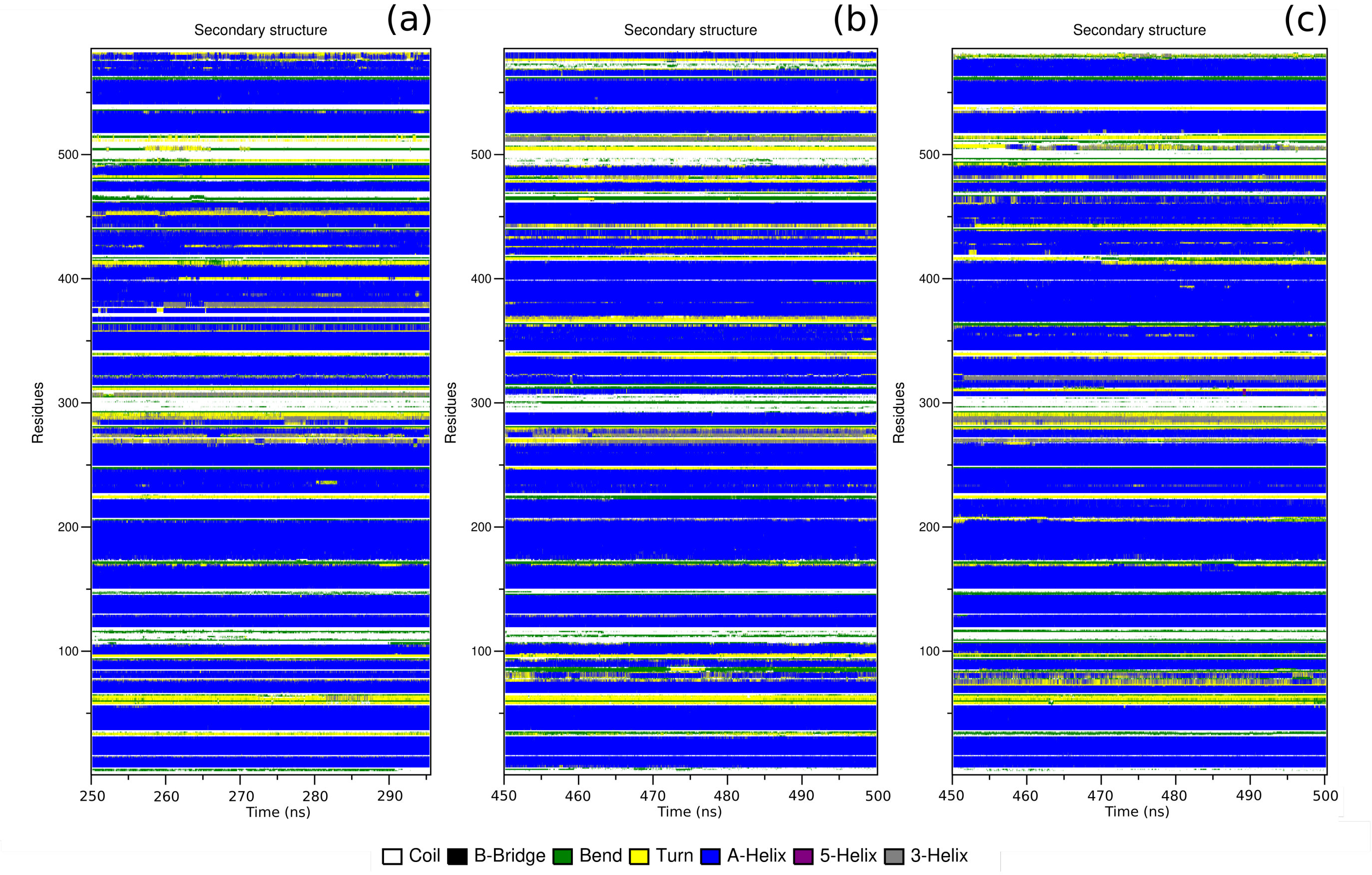


Figure 6: Secondary structure of BSA in pure water (a) and in ionic liquid (IL)/water solutions of different concentrations: (b) 1 BSA/100 IL, (c) BSA/200 IL).
